# Taxon appearance from extraction and amplification steps demonstrates the value of multiple controls in tick microbiota analysis

**DOI:** 10.1101/714030

**Authors:** Emilie Lejal, Agustín Estrada-Peña, Maud Marsot, Jean-François Cosson, Olivier Rué, Mahendra Mariadassou, Cédric Midoux, Muriel Vayssier-Taussat, Thomas Pollet

## Abstract

**Background:** The development of high throughput sequencing technologies has substantially improved analysis of bacterial community diversity, composition, and functions. Over the last decade, high throughput sequencing has been used extensively to identify the diversity and composition of tick microbial communities. However, a growing number of studies are warning about the impact of contamination brought along the different steps of the analytical process, from DNA extraction to amplification. In low biomass samples, e.g. individual tick samples, these contaminants may represent a large part of the obtained sequences, and thus generate considerable errors in downstream analyses and in the interpretation of results. Most studies of tick microbiota either do not mention the inclusion of controls during the DNA extraction or amplification steps, or consider the lack of an electrophoresis signal as an absence of contamination. In this context, we aimed to assess the proportion of contaminant sequences resulting from these steps. We analyzed the microbiota of individual *Ixodes ricinus* ticks by including several categories of controls throughout the analytical process: crushing, DNA extraction, and DNA amplification.

**Results:** Controls yielded a significant number of sequences (1,126 to 13,198 mean sequences, depending on the control category). Some operational taxonomic units (OTUs) detected in these controls belong to genera reported in previous tick microbiota studies. In this study, these OTUs accounted for 50.9% of the total number of sequences in our samples, and were considered contaminants. Contamination levels (i.e. the percentage of sequences belonging to OTUs identified as contaminants) varied with tick stage and gender: 76.3% of nymphs and 75% of males demonstrated contamination over 50%, while most females (65.7%) had rates lower than 20%. Contamination mainly corresponded to OTUs detected in crushing and DNA extraction controls, highlighting the importance of carefully controlling these steps.

**Conclusion:** Here, we showed that contaminant OTUs from extraction and amplification steps can represent more than half the total sequence yield in sequencing runs, and lead to unreliable results when characterizing tick microbial communities. We thus strongly advise the routine use of negative controls in tick microbiota studies, and more generally in studies involving low biomass samples.

## Introduction

Analysis of microbial community composition by 16S tag sequencing represents a powerful tool commonly used to evaluate the prokaryotic diversity in many ecosystems. Thanks to high-throughput sequencing approaches, it is now feasible to identify microbiota with high coverage and reveal the exceptional and complex microbial diversity of environments ranging from the wide ocean (Sogin *et al.*, 2006; Hamdan *et al.*, 2013) to small arthropods (Degli Esposti and Martinez Romero, 2017; Estrada-Peña *et al.*, 2018). With the constant improvement of sensitivity associated with its decreasing cost, 16S tag sequencing has spread to many areas, including those dealing with low biomass communities such as tick microbiota.

Despite the limitation induced by low biomass, a high number of tick microbiota studies are focusing on individual ticks (Lalzar *et al.*, 2012; Hawlena *et al.*, 2013; Rynkiewicz *et al.*, 2015; Van Treuren *et al.*, 2015; Gall *et al.*, 2016; Khoo *et al.*, 2016; Kwan *et al.*, 2017; Swei and Kwan, 2017; Estrada-Peña *et al.*, 2018) or even on individual or pooled tick organs (Andreotti *et al.*, 2011; Budachetri *et al.*, 2014; Narasimhan *et al.*, 2014; Qiu *et al.*, 2014; Clayton *et al.*, 2015; Gall *et al.*, 2016; Zolnik *et al.*, 2016; Abraham *et al.*, 2017). The results obtained from these studies have improved our knowledge of tick microbiota, while demonstrating links between its composition and several factors, such as tick stages and gender (Lalzar *et al.*, 2012; Van Treuren *et al.*, 2015; Zolnik *et al.*, 2016; Swei and Kwan, 2017), organ of origin (Zolnik *et al.*, 2016), living environment (Carpi *et al.*, 2011; Zolnik *et al.*, 2016; Estrada-Peña *et al.*, 2018), host blood meal (Swei and Kwan, 2017), or tick engorgement (Moreno *et al.*, 2006; Heise *et al.*, 2010; Narasimhan *et al.*, 2014). The presence of tick-borne pathogens has also been demonstrated to be related to differences in tick microbiota composition and diversity (Narasimhan *et al.*, 2014; Gall *et al.*, 2016; Abraham *et al.*, 2017). The continuous improvements in our knowledge, and in particular the assessment of potential links between tick-borne pathogens and particular microbiota members, require us to continue performing analyses on individual ticks, or even better, on individual tick organs.

Several potential biases, inherent to the PCR amplification (chimeric sequences) or sequencing steps (random sequencing errors), are now well known (Galan *et al.*, 2016), and pipelines have been developed to detect and remove these errors during the sequence analysis. However, several studies have begun to warn scientists about the potential distortion of results due to contamination from the extraction/amplification steps or from the environment during sample processing (Salter *et al.*, 2014; Weiss *et al.*, 2014; Narasimhan and Fikrig, 2015; Galan *et al.*, 2016; Glassing *et al.*, 2016; Knight *et al.*, 2018). This potential contamination is able to significantly modify results concerning microbe diversity and composition, particularly in the case of low biomass samples, and can have serious consequences on the interpretation of results. For example, while investigating 16s rRNA gene profiles from nasopharyngeal swabs regularly sampled from a cohort of children from 1 to 24 months, Salter *et al*. (2014) demonstrated that the apparent clustering of samples by age, observed by principal coordinate analysis, in fact corresponded to the commercial kit used to perform the DNA extraction of these samples. While studying the influence of laboratory processing on diluted samples, these authors also observed that the lower the initial biomass of samples, the higher the proportion of contaminants associated with the kits or the environment. In the context of tick microbiota studies, performing using negative controls would therefore be essential to detect potential contamination, and efficiently distinguish this contamination from genuine tick microbial communities. On the basis of an exhaustive review of the scientific literature, it is very clear that routine multiple control analyses (extraction and PCR negative controls) are not considered at their true value, and are only rarely implemented in tick microbial community analyses (Hawlena *et al.*, 2013; Gofton *et al.*, 2015; Rynkiewicz *et al.*, 2015; René-Martellet *et al.*, 2017; Sperling *et al.*, 2017; Estrada-Peña *et al.*, 2018; Thapa *et al.*, 2019). In this context, we performed a 16s rRNA gene sequencing analysis on individual ticks (females, males and nymphs) considering concomitantly several negative controls at each step of the process, prior to the sequencing run (tick crushing, DNA extraction and amplification).

## Methods

### Tick collection and sample selection

Questing *Ixodes ricinus* nymphs and adults were collected for three years by dragging (from April 2014 to May 2017) in the Sénart forest in the south of Paris. More details on the sampling location and design, and tick collection, are available in Lejal *et al*. (2019). After morphological identification, ticks were stored at -80°C until further analysis.

### Tick washing, crushing, and DNA extraction

Ticks were first washed once in ethanol 70% for 5 minutes and rinsed twice in sterile MilliQ water for 5 minutes each time. Ticks were then individually crushed in 375 µL of Dulbecco’s Modified Eagle Medium with decomplemented fetal calf serum (10%) and six steel beads using the homogenizer Precellys^®^24 Dual (Bertin, France) at 5,500 rpm for 20 seconds. DNA extraction was performed on 100 µL of tick crushing, using the NucleoSpin^®^ Tissue DNA extraction kit (Macherey-Nagel, Germany). For these previous steps, sterile tubes and filter tips were used on a dedicated area for DNA extraction. Bench and materials were decontaminated with DNA remover (Molecular BioProducts, San Diego, CA, USA) before and after each use. Each time a tick crushing or DNA extraction was performed, a crushing or extraction control was added for the corresponding step.

### DNA amplification and multiplexing

In total, 32 males, 35 females, and 557 nymphs were subjected to sequencing analysis. DNA amplifications were performed on the V4 region of the 16s rRNA gene using the primer pair used by Galan *et al*. (2016) (16S-V4F: 5’-GTGCCAGCMGCCGCGGTAA-3’ and 16S-V4R: 5’-GGACTACHVGGGTWTCTAATCC-3’), producing a 251 bp amplicon. This primer pair was chosen for its ability to specifically amplify prokaryotic DNA (86% or 95% of referenced bacterial DNA in the Silva database, and 52.3% or 93% of Archaeal DNA, when considering 0 or 1 mismatch, respectively). This characteristic is crucial in the context of tick microbiota analysis, due to the high amount of tick DNA compared to prokaryotic DNA. An 8 bp-index was added to the forward and reverse primers. The individual amplification of each sample was performed using a combination of 22 and 32 index-tagged forward and reverse primers, allowing for amplification and multiplexing of a maximum of 704 samples that can be loaded together on an Illumina MiSeq flow cell. Amplification and multiplexing steps were performed under a sterile hood, decontaminated before and after use with DNA remover and UVs, using tube strips with individual caps to prevent cross-contamination. All the PCR amplifications were carried out using the Phusion^®^ High-Fidelity DNA Polymerase amplification kit (Thermo Scientific, Lithuania). For each sample, 5 µL of DNA extract were amplified in a 50 µL final reaction volume, containing 1X Phusion HF buffer, 0.2 µM of dNTPs, 0.2 U/mL of Phusion DNA polymerase, and 0.35 µM of forward and reverse primer. The following thermal cycling procedure was used: initial denaturation at 98°C for 30 s, 35 cycles of denaturation at 98°C for 10 s, annealing at 55°C for 30 s, followed by extension at 72°C for 30 s. The final extension was carried out at 72°C for 10 minutes. Amplification step was performed on a peqSTAR 2x thermocycler (PEQLAB, Germany). For each PCR run, negative controls were performed by using the reaction mixture without template, but using ultrapure water (Invitrogen, United Kingdom). PCR products were checked on 1.5% agarose gels.

### DNA purification, quantification, and pooling

Amplicons were cleaned and quantified using a SequalPrep normalization plate kit (Invitrogen, USA), and a Qubit dsDNA High Sensitivity Assay kit (Invitrogen, USA), respectively. Amplicons and controls were pooled at equimolar concentrations and sent to the sequencing platform (GenoScreen, France).

### Controls

In total, 45 negative controls were included and analyzed using the same process as for tick samples (**Figure 1**).

**Figure 1.**
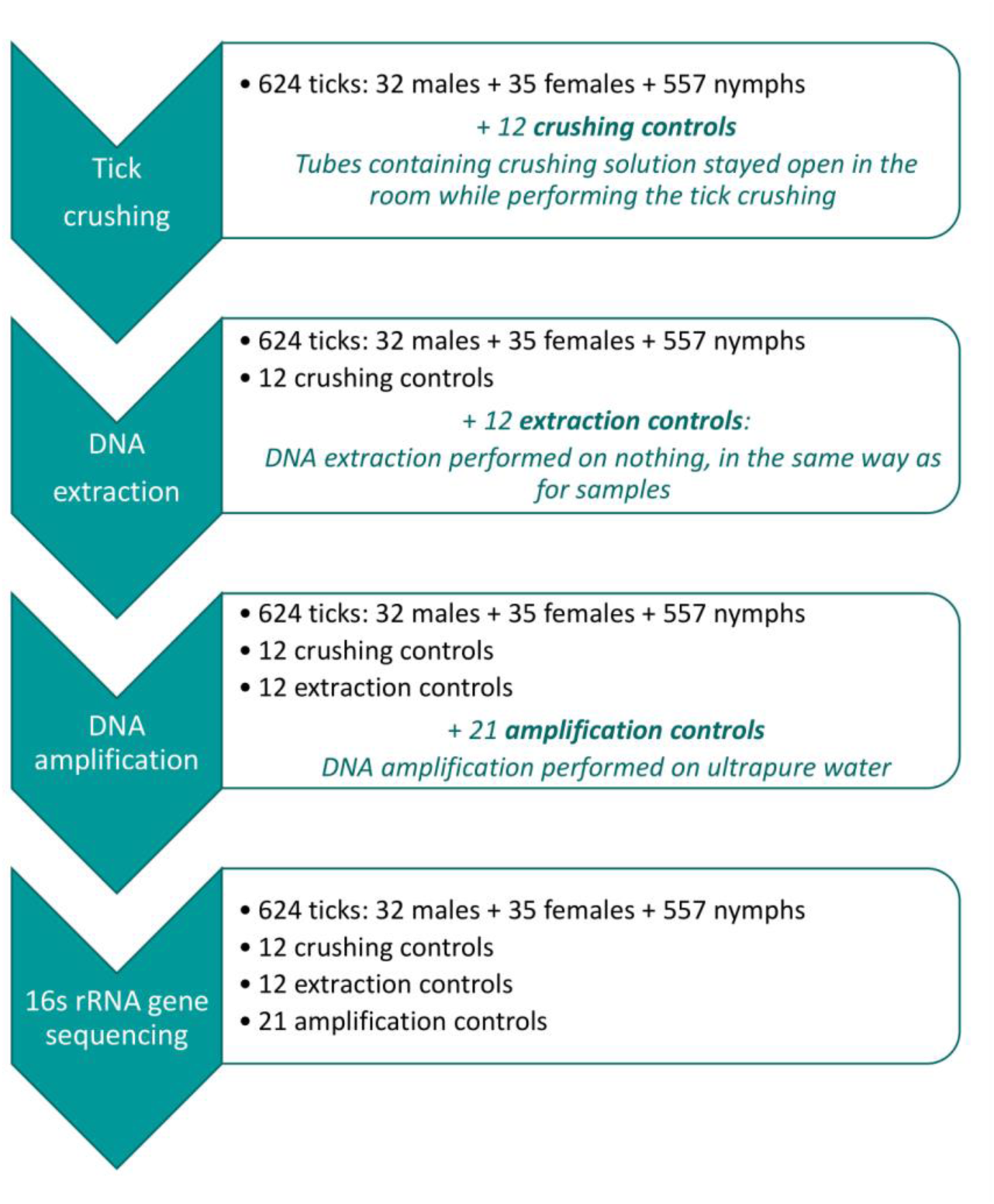
Analyzing process for samples and added controls.

#### 12 Crushing controls (CCs)

Tube containing crushing solution, remaining open in the room while putting ticks inside other tubes between the cleaning and crushing steps. DNA extraction and amplification were then performed on the crushing control in the same conditions as for any other sample.

#### 12 DNA extraction controls (ECs)

For these controls, the different DNA extraction steps were performed under the same conditions as for any other samples, but without adding sample. DNA amplification was then performed on the extraction control in the same conditions as for any other sample.

#### 21 DNA amplification controls (ACs)

The DNA amplification step was performed on ultrapure water (Invitrogen, United Kingdom), in the same conditions as for any other sample.

### Sequencing and data processing

The equimolar mix was concentrated and sequenced by GenoScreen (Lille, France) using MiSeq Illumina 2 × 250 bp chemistry. Quality control checks on raw data were performed using FastQC v0.11.3 (Andrews, 2010), and no problems were detected. First, pairs of amplicons were merged using Pear v0.9.11 (Zhang *et al.*, 2014) with a minimum overlap of 20 bp, a minimum assembly length of 100 bp, a maximum assembly length of 480 bp, and an e-value of 0.01. Then, the sequencing orientation of extended sequences was checked by using cutadapt v1.12 (Martin, 2011), with primers in forward and reverse strands (without removing them). Sequences with a reversed orientation were reverse-complemented to obtain a set of sequences with the same orientation (5’ to 3’). The sequence analysis was performed using FROGS (Escudié *et al.*, 2018). Primers were removed using cutadapt and resulting sequences were dereplicated. Operational taxonomic units (OTUs) were built using Swarm v2.1.12 (Mahé *et al.*, 2015) with parameter d = 3, and chimeras were filtered using vsearch v1.4 (Rognes *et al.*, 2016) in a de novo mode. Finally, OTUs with low abundance (< 0.005% of the total sequence count) were filtered out, resulting in 608 OTUs (corresponding to 2,835,506 sequences). OTUs were affiliated by blasting cluster seed sequences against the Silva database v132 (Quast *et al.*, 2013). OTUs corresponding to mitochondrial/chloroplast DNA or not affiliated to any domain were removed, leading to a dataset composed of 513 OTUs and 2,726,287 sequences.

### Data processing to distinguish contaminant versus tick microbiota OTUs

The control composition was used to distinguish tick microbial OTUs from contaminants. All the OTUs corresponding to more than 1% of the total number of sequences obtained in each category of control were identified as main contaminants. However, due to possible errors in sequence assignment occurring during sequencing due to the formation of mixed clusters on the flow cell (Kircher *et al.*, 2012), negative controls can also contain a few sequences of tick microbial OTUs, complicating contaminant identification under this 1% threshold. Therefore, for each OTU, the number of sequences expected to spread to other samples due to false assignment (i.e. expected number) was determined by calculating 0.02% of the total number of each OTU in the dataset, corresponding to a false assignment rate estimated previously by Galan *et al*. (2016). This expected number was compared to the maximum number of obtained sequences in controls (i.e. maximum number). As this rate was not estimated from our dataset, we decided to allow variability around this threshold, while calculating a 99% confidence interval around the expected number of 0.02%. OTUs whose maximum sequence number was above the upper threshold were considered contaminants. Importantly, cross-contamination from samples to controls was considered negligible, as no negative controls presented a higher maximum sequence number than the expected number for an OTU belonging to the most abundant and characteristic genus of tick microbiota in our dataset (Midichloria), which is also the 3^rd^ most abundant OTU in the dataset (266,516 sequences).

### Statistical analyses

The percentage of samples presenting a contamination rate higher than 50% was compared between samples categories (nymphs, males and females) according to a Chi^2^ statistical test.

Alpha diversity was estimated before and after sample cleaning. Samples containing fewer than 500 sequences after cleaning were removed from the analysis, even if they contained more than 500 sequences before cleaning. Several measures of alpha diversity were calculated. Basically, the observed number of OTUs (species richness) was used as well as the Faith Phylogenetic Diversity (PD) index, corresponding to the total branch length of the subtree spanned by the community, and also two indexes taking into account the number of sequences: the Shannon index (more influenced by rare OTUs) and Simpson index (more influenced by dominant OTUs). Phylogenetic diversity was estimated using the Picante package (Kembel *et al.*, 2010), while other measures were assessed thanks to Easy 16S (http://genome.jouy.inra.fr/shiny/easy16S/), based on the phyloseq package (McMurdie and Holmes, 2013). Alpha diversity measure comparison was performed between cleaned and not cleaned groups, as well as between stages, using non-parametric statistical tests (Wilcoxon and Kruskal–Wallis, respectively). Statistical computations were performed in R 3.6.0 (R Core Team, 2018).

## Results

Nymphs, males and females presented a mean number of sequences per sample of 3,721, 2,689 and 5,316, respectively. Concerning controls, a mean number of 13,198, 11,235 and 1,126 sequences per sample was obtained for crushing, extraction and amplification controls (CCs, ECs and ACs), respectively.

The most abundant OTUs, with relative abundance higher than 1% in at least one category of samples or controls, are represented in **Figure 2**. All other OTUs, i.e. those with relative abundance lower than 1% in all categories were grouped together in the “Others” category.

**Figure 2.**
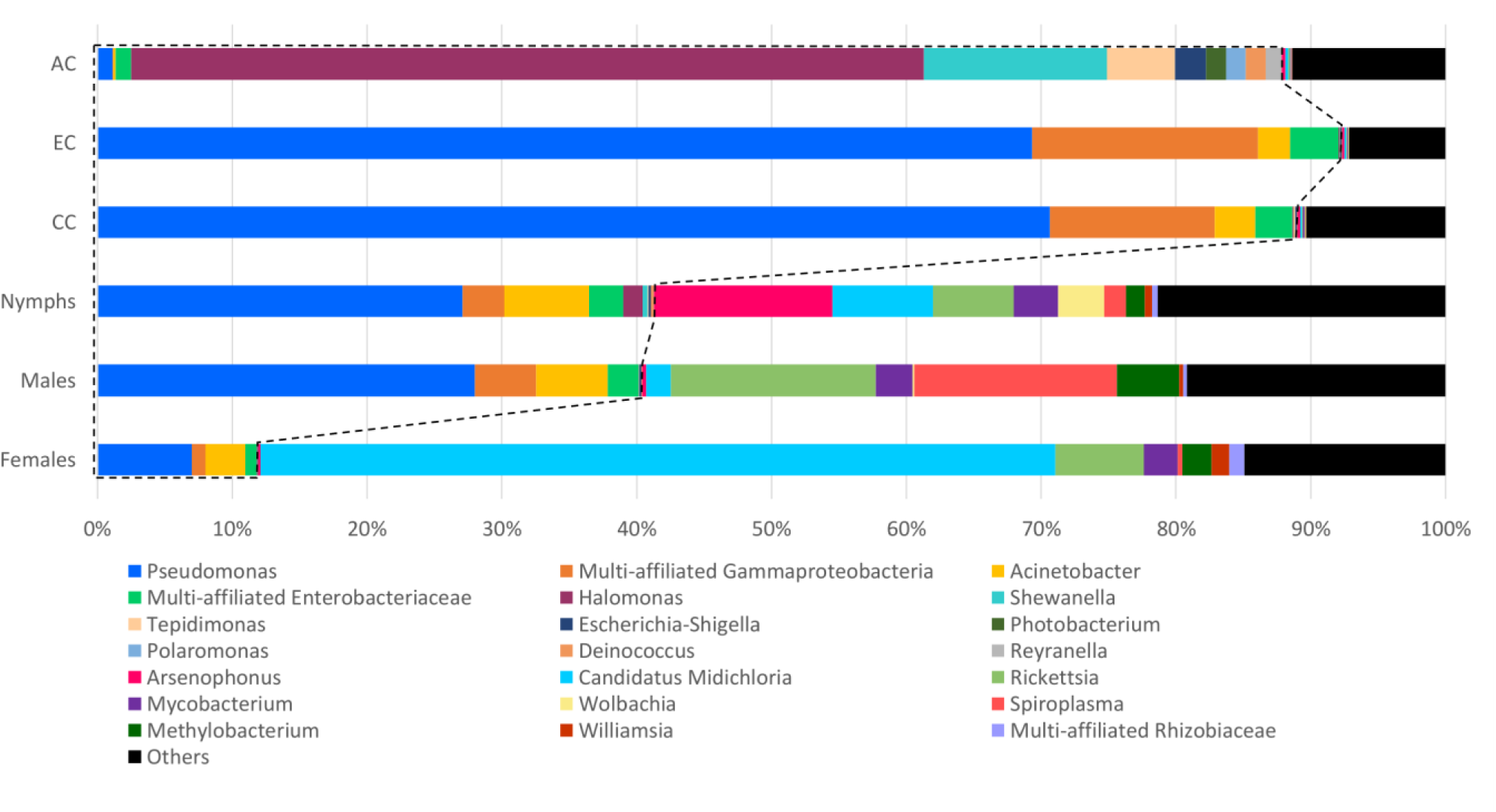
Comparison of the 16S rRNA gene sequencing taxonomic assignments between negative controls and samples. Percentages are represented for the OTUs with proportions corresponding to at least 1% of the total number of sequences assigned to a category of samples or controls. Those corresponding to less than 1% are assigned as “Others”. OTUs belonging to the same genus or family (if multi-affiliated at the scale of genus) were assigned under the same name. One OTU was only identified until the class level (multi-affiliated Gammaproteobacteria).

The examination of the at-least-1% OTU composition of the control (CCs, ECs, ACs) allowed to identify OTUs belonging to the genera: *Pseudomonas, Acinetobacter, Halomonas, Shewanella, Tepidimonas, Esherichia-Shigella, Photobacterium, Polaromonas, Deinococcus* and *Reyranella*, as well as two OTUs belonging to the Enterobacteriaceae family but multi-affiliated at the genus level, and one OTU only affiliated to Gammaproteobacteria at the Class level (**Figure 2**). OTU composition was contrasted across the control types. While the same OTUs were identified in CCs and ECs and belonged to *Pseudomonas* [70.7% and 69.3%, respectively], multi-affiliated Gammaproteobacteria [12.2% and 16.7%, respectively], *Acinetobacter* [3.1% and 2.4%, respectively], and multi-affiliated Enterobacteriaceae [2.7% and 3.6%, respectively]), the microbial community identified in AC controls was dominated by OTUs belonging to *Halomonas* (58.8%), *Shewanella* (13.6%), *Tepidimonas* (5.1%), *Escherichia-Shigella* (2.3%), *Photobacterium* (1.5%), *Polaromonas* (1.5%), *Deinococcus* (1.5%), *Reyranella* (1.2%) and multi-affiliated Enterobacteriaceae (1.2%). These OTUs represented 38.9% of the total number of sequences coming from the tick samples. Focusing on the sample category, these OTUs correspond to 11.9%, 40.4% and 41.3% of sequences detected in females, males and nymphs, respectively (**Figure 2**). These OTUs were therefore considered to be the main contaminants of the dataset. It is interesting to note that most of these OTUs correspond to those detected in both EC and CC controls. In fact, only 2.8% of sequences detected in the different categories of samples (nymphs, females and males) correspond to OTUs detected in the AC controls and identified as main contaminants.

A more in-depth identification of contaminants, including those corresponding to less than 1% of the total number of sequences was performed, allowing us to identify 160 OTUs as contaminants among the 513 initially identified OTUs in the dataset. This identification increased the proportions of identified contaminants in the samples, which reach 50.9% of the total sequence count from the tick samples, and 20.1%, 51.2% and 53.7% of sequences detected in females, males and nymphs, respectively. Furthermore, after these cleaning steps, 201 samples contained less than 500 sequences compared to 8 before cleaning. For 76.6% of nymph and 75% of male samples, contaminant OTUs represented more than 50% of the total sequence count (**Figure 3**). These sample contamination rates are significantly higher than those obtained for females (p < 0.05 according to a Chi^2^ test), for which most of the samples (65.7%) exhibited a contamination rate below 20%.

**Figure 3.**
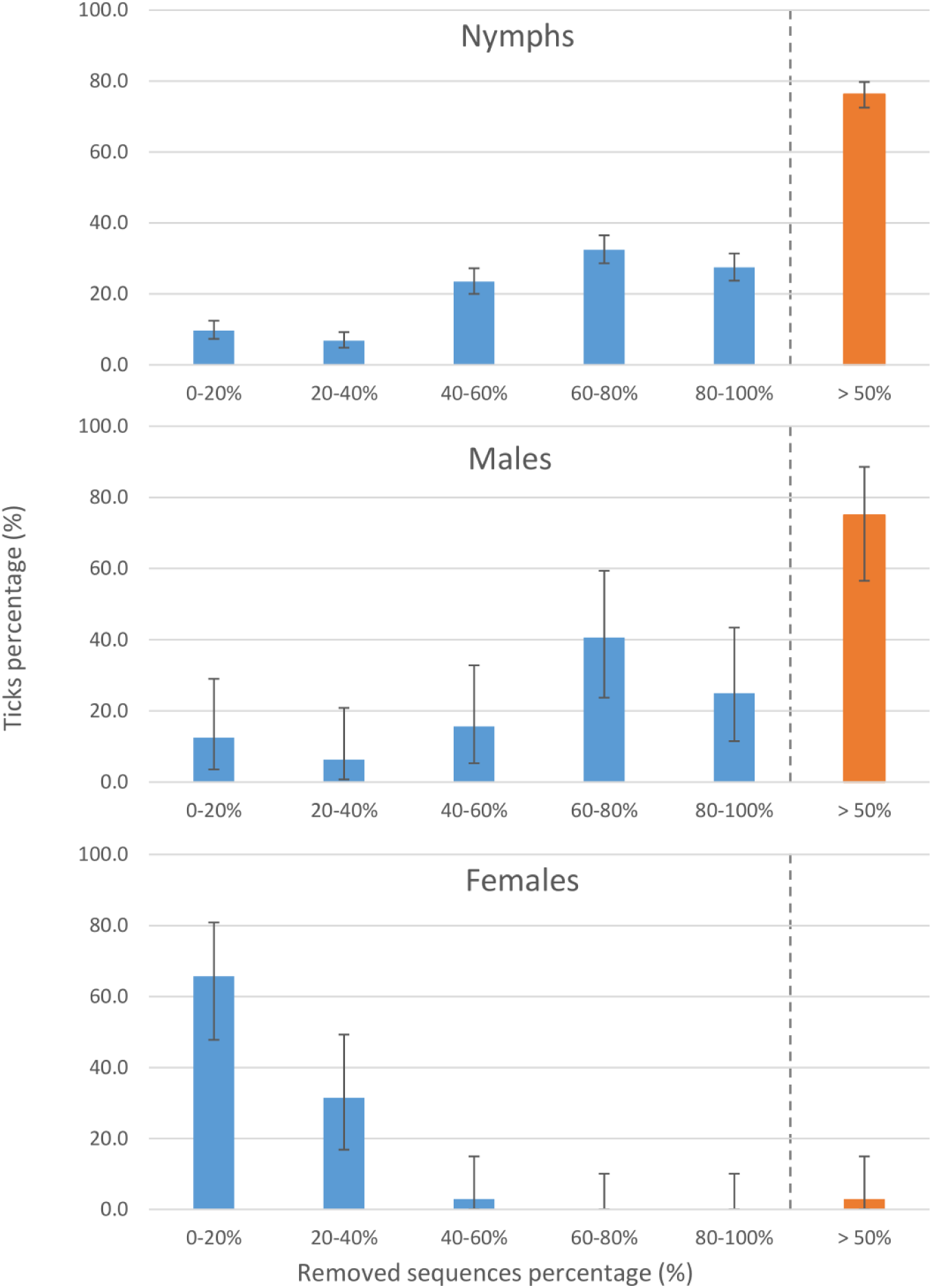
Impact of contamination on the final number of sequences per sample according to the tick stage/gender.

The impact of this contamination on diversity, according to the sample category was estimated (**Figure 4**). As expected, for all the studied diversity measures, cleaning led to a decrease in diversity. Concerning nymphs and females, this decrease was significant for all the tested diversity indexes, while for males, it was only significant for the observed and phylogenetic diversity measures, suggesting that for males, contamination concerned mostly low-abundance taxa. The impact of this cleaning on alpha diversity comparison between tick stages was been studied. Without cleaning, significant differences in alpha diversity were found between tick stages for observed, Shannon and Simpson measures. Concerning cleaned data, significant differences were still observed between stages when using Shannon and Simpson indexes, but disappeared when using the observed diversity measure.

**Figure 4.**
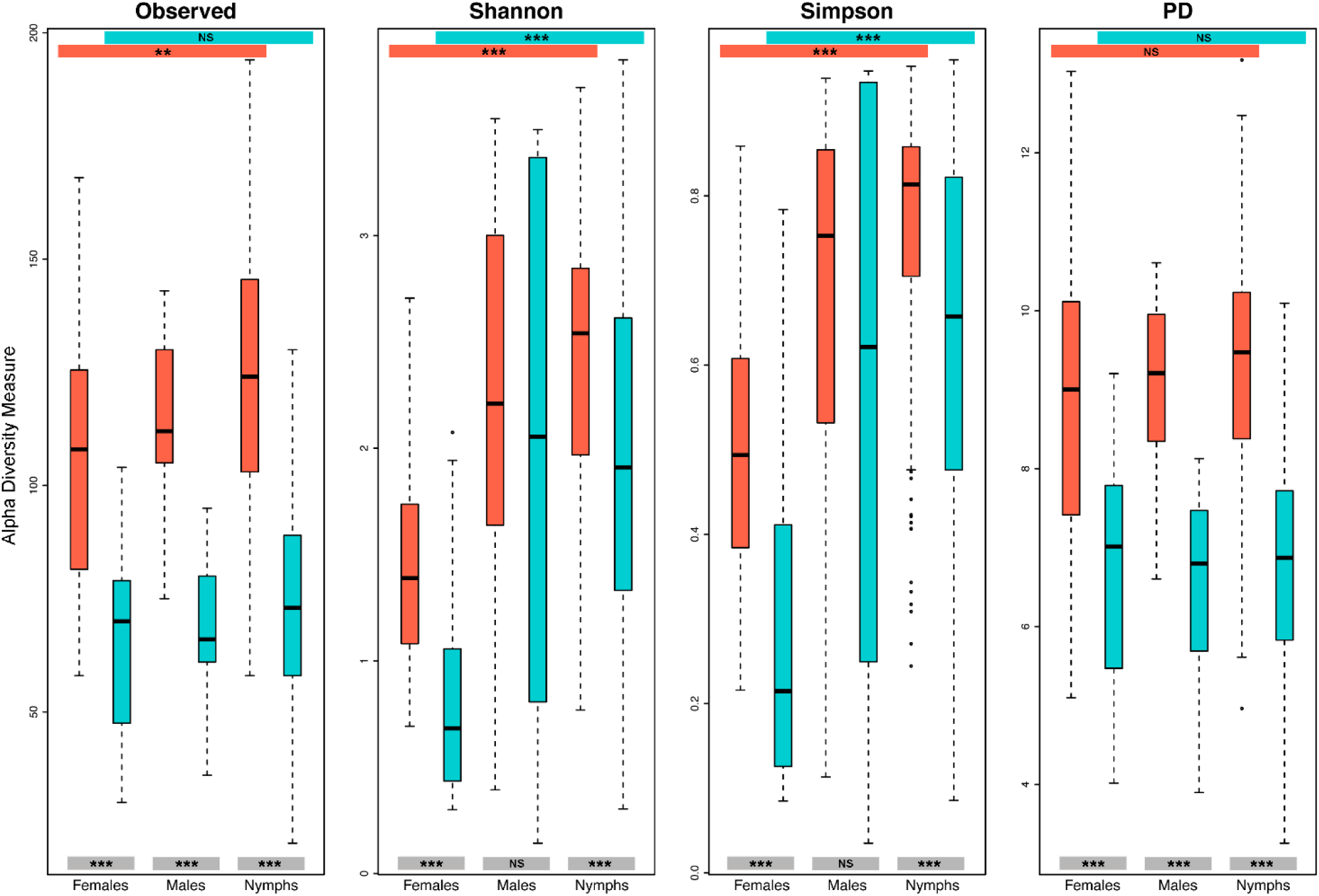
Impact of contamination on alpha diversity estimations according to the tick stages. Alpha diversity has been estimated according to the observed number of OTUs, the Shannon and Simpson indexes and the Faith’s phylogenetic diversity (PD), for each stage on cleaned (turquoise) and not cleaned (orange-red) datasets. Results of the statistical test comparing alpha diversity measurement between stages, as well as between the cleaned and uncleaned groups, are represented on coloured bands at both the top and the bottom of the frame respectively: NS= Not Significant, * * * = p < 0.001, * * = p < 0.01.

## Discussion

In this study, we sequenced the microbiota of 624 individual *I. ricinus* ticks including several categories of negative controls to identify and quantify the contaminating sequences, in particular those produced during the extraction or amplification steps. The performance of multiple negative controls allowed us to perform in-depth identification of contaminant OTUs. These OTUs represented 50.9% of the total sequence yield from tick samples, and had a significant impact on microbial diversity estimation. This shows the importance of contamination potentially arising from processing steps, and demonstrates that its consequences on data analyses can be significant and should not be neglected.

A high number of sequences in each category of controls was detected, particularly in CCs and ECs, in which the mean number of sequences was even higher than the mean sequence number detected in samples. Importantly, this higher number might be partly overestimated, first due to the presence of tick DNA during the PCR that may have inhibited the reaction on the 16s rRNA gene in samples while not in controls, and second during the measure of DNA concentrations that may have led to a higher estimation of concentrations in samples compared to controls, before the pooling step. Despite this, the high number of sequences in controls raises the issue of contaminant presence, while performing each step of the sample processing.

Each processing step leads to a different contamination pattern. Most of the main contaminants do not seem to originate from the DNA amplification step. Furthermore, as the composition of crushing and extraction controls was the same, while the reagents used were different and crushing control also underwent the extraction step, the contaminant sequences most probably came from the DNA extraction step, rather than from the crushing step. The key role of the extraction step in the contamination of low biomass samples has previously been demonstrated by Salter *et al*. (2014). Furthermore, three of the four main contaminants identified in our analysis, corresponding to *Pseudomonas, Acinetobacter*, and multi-affiliated Enterobacteriaceae, belong to families and genera identified by these authors as contaminants coming from DNA extraction kits. Our observations differed from those obtain by Galan *et al*. (2016), who mainly detected contaminants coming from the DNA amplification step, while this step represented only 2% of the main contaminants detected in our samples. In their study of kit contaminants, Salter *et al*. (2014) observed that the extent of contamination arising from DNA extraction was different depending on the kit brand used to perform this step. While using the less contaminating kit, they extracted DNA from several 10-fold cascade dilutions of a pure suspension of *Salmonella bongori*, and subsequently assessed 16s rRNA gene concentrations *via* q-PCR. They demonstrated a linear relationship between the 16s rRNA gene copy number and the dilution fold only for the first dilution points, after which levels started to stabilize and to reach a threshold of 10^5^ mean copy number/mL. Consequently, the higher rate of extraction, rather than amplification contaminants in samples, could be due either to variations of contaminating load between our extraction and amplification kits, that are not sourced from the same company, or to the presence of a basal level of extraction contaminant bacterial DNA, potentially limiting the contamination arising from the next step (amplification).

Interestingly, we showed that in terms of contamination rates, the influence of contaminant presence was significantly higher in nymph and male samples compared to female ones. Because low biomass samples were shown to be particularly sensitive to contamination (Salter *et al.*, 2014), we might suppose that the amount of bacterial DNA would be lower in males and nymphs, explaining the higher presence of contaminants in these samples. It would be useful to develop epifluorescence or flow cytometry protocols adapted to tick samples, to assess whether the amount of bacteria composing the female microbiota is effectively higher than in the male and nymph. Additionally, previous studies dealing with tick microbes have identified higher microbiota diversity in males and nymphs compared to females, while investigating richness and PD (Van Treuren *et al.*, 2015; Zolnik *et al.*, 2016) as well as Shannon and Simpson indexes (Zolnik *et al.*, 2016). However, no negative controls and data cleaning were reported in these studies, and they could therefore result from contamination. In the present study, we determined that the presence of contaminants can lead to an overestimation of microbial diversity in all the tick stages, and that this is the case for several diversity indexes, especially richness and PD estimations, which are the measures most affected by the cleaning process. Furthermore, the impact of removing contaminants on diversity comparisons between stages was estimated and showed that this cleaning step can influence the final results, and particularly in our case, concerning the observed diversity estimation. Our results therefore call for caution in attempts to draw conclusion about tick stage diversity without considering negative controls.

As already mentioned by Narasimhan and Fikrig (2015), the understanding of arthropod microbial communities in the context of arthropod survival and pathogen transmission may open new routes in arthropod-borne pathogen control strategies. In this context, many studies have assessed tick microbiota composition in individual ticks or even in tick organs (Andreotti *et al.*, 2011; Lalzar *et al.*, 2012; Hawlena *et al.*, 2013; Budachetri *et al.*, 2014; Narasimhan *et al.*, 2014; Qiu *et al.*, 2014; Clayton *et al.*, 2015; Rynkiewicz *et al.*, 2015; Van Treuren *et al.*, 2015; Gall *et al.*, 2016; Khoo *et al.*, 2016; Zolnik *et al.*, 2016; Abraham *et al.*, 2017; Kwan *et al.*, 2017; Swei and Kwan, 2017; Estrada-Peña *et al.*, 2018). However, very few reported the use of negative controls or mentioned the identification and removal of subsequently identified contaminants (Hawlena *et al.*, 2013; Rynkiewicz *et al.*, 2015; Estrada-Peña *et al.*, 2018), thus obscuring the risk that identified micro-organisms as members of the microbiota could correspond to contaminants. Nonetheless, the bias generated by the presence of contaminants can have a considerable influence on the characterization of tick microbiota composition and structure. Here, we showed that contaminants represented almost 50 % of the total sequence yield of tick samples. Furthermore, several contaminants we identified belong to genera (especially *Pseudomonas* and *Acinetobacter*) previously reported in several tick microbiota studies (reviewed by Narasimhan and Fikrig, 2015). This does not necessarily mean that these OTUs are contaminants, as several OTUs belonging to the same genera may correspond either to tick microbiota members or to contaminants. However, this points out the lack of reliability concerning this crucial point in the understanding of tick microbiota composition, and the need to perform negative controls to clean the data set and to rule out the hypothesis of potential contamination.

## Conclusions

Because of their ease of use and the fact that they employ safer chemicals than those in conventional methods (i.e. phenol chloroform extraction), the use of commercial DNA extraction kits has become standard, particularly in the context of studies dealing with the identification of tick microbial communities. However, the issue of contaminants coming from DNA extraction kits has recently been raised for microbiota studies dealing with low biomass samples, such as blood, intestinal tissue, mucosal tissue, or nasopharyngeal samples. Our study adds further evidence supporting these important concerns in the field of tick microbiota. Here, our results suggest that many negative controls should be performed at each step of sample processing prior to performing microbiota analyses, allowing us to efficiently identify a potentially serious bias in the study of the actual tick microbial community. While more and more studies try to identify tick microbiota to *in fine* potentially propose hypothesis for the development of new tick-borne pathogen control strategies, we strongly advise the routine use of negative controls in microbiota studies, particularly in the context of low biomass samples like ticks or other small arthropods. These procedures should be used routinely in order to obtain the most reliable results possible.

## List of abbreviations

OTU: Operational Taxonomic Unit
*I. ricinus*: *Ixodes ricinus*
CC: Crushing Control
EC: Extraction Control
AC: Amplification Control

## Author Contributions

Conceived and designed the experiments: TP, MVT, JFC, EL. Performed the experiments: EL. Analysed the data: EL, AE, TP, MM, MM, CM, OR. Wrote the paper: EL, AE, MM, JFC, OR, MM, CM, MVT, TP. All the authors read and approved the final manuscript.

## Acknowledgments

The authors would like to thanks Sabine Delannoy and the IdentyPath genomic platform that allowed us to perform part of the experiments in their laboratory. We also thank Maxime Galan for providing index sequences and stimulating discussions. Thank you as well to Ladislav Šimo for his precious help in figure improvement.

